# MoBIE: A Fiji plugin for sharing and exploration of multi-modal cloud-hosted big image data

**DOI:** 10.1101/2022.05.27.493763

**Authors:** Constantin Pape, Kimberly Meechan, Ekaterina Moreva, Martin Schorb, Nicolas Chiaruttini, Valentyna Zinchenko, Hernando Vergara, Giulia Mizzon, Josh Moore, Detlev Arendt, Anna Kreshuk, Yannick Schwab, Christian Tischer

## Abstract

Facing the challenge of exploring and sharing multi-terabyte, multi-modal and multi-scale image data of heterogeneous dimensionality, we developed MoBIE, a Fiji plugin that provides rich visualization features to enable browsing data from numerous biomedical applications on a standard laptop computer. MoBIE also supports segmentations, associated measurements and annotations. Users can configure complex views of datasets, share them with collaborators, and use them for interactive figure panels. The MoBIE plugin also offers a convenient interface for converting data into compatible data formats; an additional Python library facilitates managing diverse MoBIE projects.

## Main

Modern microscopy produces massive image datasets that enable detailed multi-scale analysis and can combine several modalities, for example in correlative microscopy. Visualizing, exploring and sharing such data are significant challenges both during the execution of a research project, and after publication to enable open access and re-use^1^. To this end we have developed MoBIE, a Fiji^2^ plugin for **M**ultiM**o**dal **B**ig **I**mage Data Sharing and **E**xploration. It supports visualization of multi-scale data of heterogeneous dimensionality (i.e. combined 2D, 3D or 4D data), several-TB-size image data, as well as the exploration of image segmentations and corresponding measurements and annotations. MoBIE uses next-generation image file formats, such as OME-Zarr^3^ or N5^4^, that enable accessing multi-scale data on local storage or via the cloud, permitting the transparent sharing and publication of data without the need to run a web-service. In addition, MoBIE allows users to easily configure and share fully reproducible complex *views* of their data. MoBIE is implemented using existing tools wherever possible, in particular BigDataViewer^5^ (BDV) and the ImgLib2^6^ and N5-java^4^ libraries. MoBIE has enabled integration of multiple modalities and open access for data from different domains of the Life Sciences. Fig. 1 shows examples from developmental biology^7^ (panel a), correlative microscopy (panel d) and high-throughput screening microscopy^8^ (panel b) and Suppl. Fig. 1 shows further examples. A video demonstrating MoBIE’s main functionality is available at https://youtu.be/oXOXkWyIIOk and its full documentation at https://mobie.github.io/.

**Figure 1:**
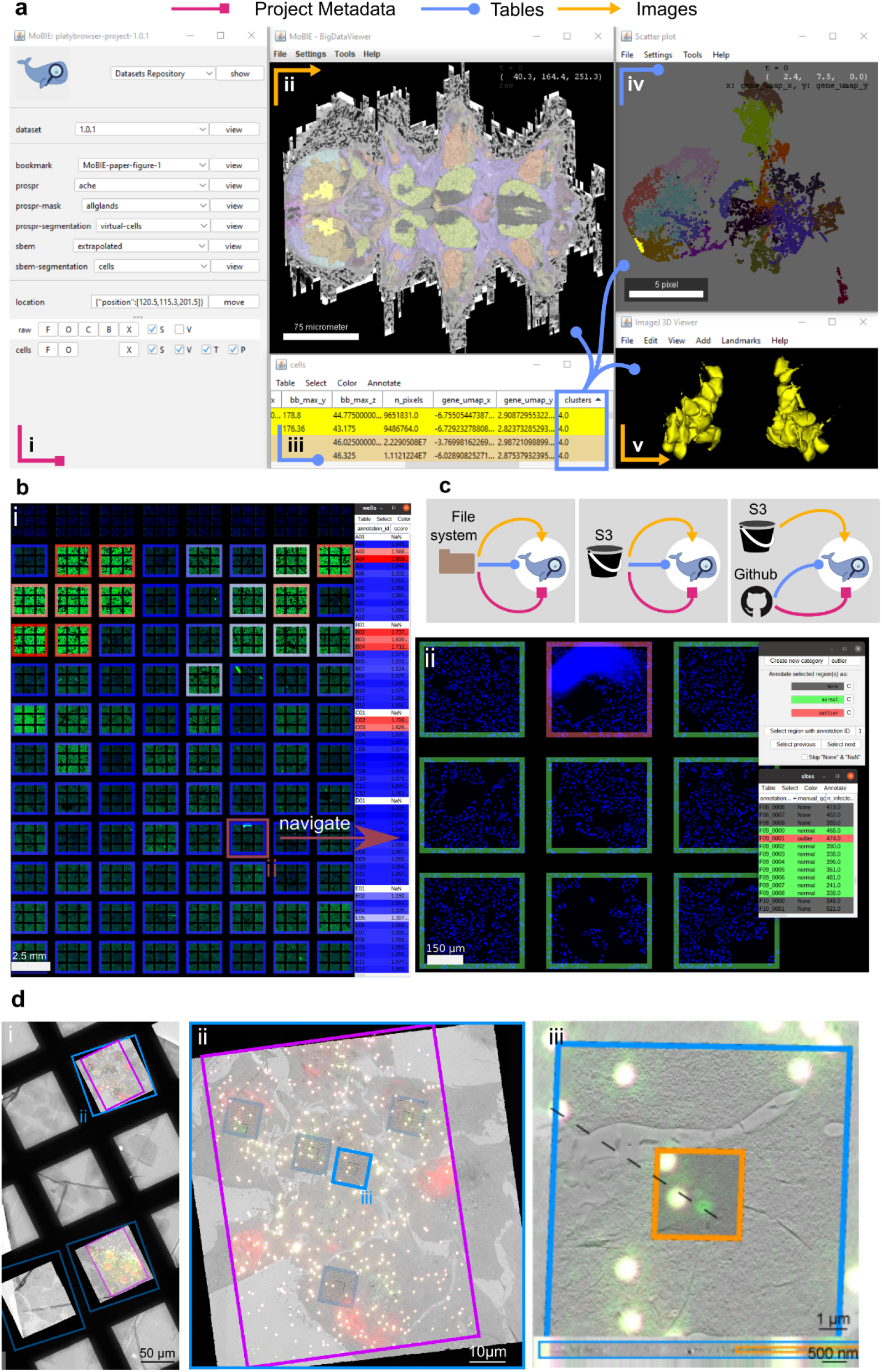
**a** - overview of the MoBIE user interface displaying the *Platynereis* atlas^7^. **i** - MoBIE controls, *view* selector corresponding to this panel is highlighted in yellow. **ii** - Main viewer (BigDataViewer) displaying a cross-section through the electron microscopy (EM) data and cell segmentation, coloured by gene expression clusters. **iii** - interactive table, showing data derived from the cell segmentation. **iv** - interactive scatter plot, showing the gene expression space coloured by gene cluster assignments. **v** - 3D viewer, displaying a subset of cells from genetic cluster 4 (same as highlighted in yellow in ii, iii and iv). **b** - High-throughput screening microscopy data^8^ shown in MoBIE. **i** - sites and wells arranged in plate layout using a nested grid transformation; well outlines are covered by per-well score measurements. **ii** - zoom-in on a single well from i) with active user annotation mode, used here to add a table for manual image quality control. **c** - the combinations of sources for metadata, tables and image data supported by MoBIE. **d** - correlative light electron microscopy data. **i** - low resolution overview EM image (2D) with registered light microscopy (LM) data (2D), high-resolution EM (2D) and EM tomograms (3D). **ii** - zoom-in to one of the positions from i), with tomograms highlighted by outlines with MoBIE region displays. **iii** - zoom-in to one of the tomograms from ii), showing the low- and high-resolution tomogram, overlaid with occluding blending modes. The lower part of the panel shows the same view across the z-axis, with the natively 3D tomograms and the 2D LM and EM data. The panels from a), b) and d) can be recreated by selecting the “MoBIE: Figure 1a” etc. items from MoBIE’s “Open MoBIE Published View…” command.

MoBIE extends BDV’s functionality to efficiently stream data from remote data storage, conveniently manage an arbitrary number of images and adds a menu to select and control *sources, views* (see below) and navigation. It inherits and extends BDV’s functionality for on-the-fly data transformations, which enable registering multi-modal data without the need to resample the (potentially TB-sized) data. Transformations can be determined with an external tool^9^ or obtained in MoBIE itself, either manually or by landmark based registration provided through BigWarp^10^ integration. MoBIE also supports joint visualization of two- and three-dimensional (time-lapse) data and different image blending modes for multi-modal data support (Fig. 1d). We have further added interactive display of segmentation results with support for different colormaps, 3D object rendering and display of tabular data associated with the individual segments, which enables fast visual inspection of derived measurements, e.g. cell sizes or gene expression levels. Tables can also be associated with regions, for example high-throughput screening microscopy data arranged on a plate (Fig. 1b). The tables can be used for coloring, searched, sorted and used to generate scatter plots. The objects in the image, the corresponding table row and the point in the scatter plot are linked, so that one can select and navigate all data representations in synchronization. We have also implemented an annotation mode that can be used to add table columns with user observations. The full MoBIE state can be captured as a *view*, which can be shared with collaborators or be used to create interactive figure panels; panels a, b and d of Fig. 1 are created with MoBIE and available as interactive *views*. See Suppl. Note 1 for a detailed description of all MoBIE features.

To access data with MoBIE, it must be organized according to the *MoBIE project specification*, see also Suppl. Note 1. A *project* consists of *datasets* that contain all data that can be loaded in a single MoBIE instance: image data, stored as OME-Zarr or N5 files, tabular data stored as tab separated value (tsv) files and a JSON file specifying the dataset layout. The *projec*t can be stored either locally or on object storage, metadata and tables can also be hosted on github for version control. The Fiji plugin can access projects from these configurations, which enables using MoBIE to access and share data throughout the full life-cycle of a project: when the data is only accessible to a single researcher or institution, accessible to a research collaboration or published and freely accessible, see Fig. 1b. MoBIE projects can be created either using the Fiji plugin, providing a convenient graphical user interface, or a python library, providing full flexibility and support for large data.

Overall, MoBIE enables scientists to seamlessly access, explore and share their massive microscopy data through all stages of a project in a convenient Fiji plugin. We are working with online archives to deploy MoBIE to support on-demand access to existing scientific image data and have already implemented a prototype with the BioImage Archive^11^. Furthermore, we are convinced that many of MoBIE’s features are of general use and aim to contribute its functionality into “upstream” core components such as BDV. To promote cross-tool accessibility, we are planning to standardize some of the MoBIE specifications, for example, transformations, views and tables, within OME-Zarr. This integration will enable other tools, for example Viv^12^, to access the same data and *views* as MoBIE.

## Acknowledgements

C.P. has received funding from the European Union’s Horizon 2020 research and innovation programme under grant agreement number 824087. C.T. acknowledges support by grant number 2020-225265 from the Chan Zuckerberg Initiative DAF, an advised fund of Silicon Valley Community Foundation. J.M. also received funding from the Chan Zuckerberg Initiative DAF for work on OME-NGFF by grant number 2019-207272 and on Zarr by grant numbers 2019-207338 and 2021-237467. D.A. is supported by a grant from the 728 European Research Council (NeuralCellTypeEvo 788921).

We would like to thank Tobias Pietzsch, John Bogovic, and Stephan Saalfeld for invaluable help with ImgLib2 and BigDataViewer code development; the EMBL IT department, in particular Josep Moscardo, for supporting the S3 infrastructure; Siân Culley for inventing the name MoBIE; Gemma Estrada Girona for drawing the MoBIE icon; Jonas Hartmann for providing his zebrafish data; Adrian Wolny for providing his plant root data and Clarisse Uwizeye and Johan Decelle for providing their plankton data.

## Contributions

C.P., K.M, V.Z., H.V., D.A, A.K., Y.S. and C.T have conceptualized the work. C.P, K.M, E.M., M.S., N.C., J.M. and C.T. have implemented the software. M.S. and G.M. have acquired new data. C.P., K.M, M.S., G.M. and C.T. have converted raw data into MoBIE projects. C.P., K.M., E.M., M.S. and C.T. have drafted the manuscript. All authors have provided comments on the manuscript.

## Methods

Here, we give an overview of the usage, features and implementation of MoBIE, and also compare to existing tools for multi-modal and big image data visualization. We first compare to other tools, then briefly explain how to use the MoBIE Fiji Plugin, describe the core features of the plugin, give an overview of its implementation, explain the data layout of MoBIE projects, describe how to create MoBIE projects, give a summary of our additions to the BDV/Imglib2 ecosystem, list several example problems and finally introduce the proof of concept implementations for accessing data from public deposition sites via MoBIE.

Several videos demonstrating MoBIE’s functionality are available at https://www.youtube.com/channel/UCtRtv0JkEEW5zLFO7d_nglg, the full user documentation is available at https://mobie.github.io/.

For questions or issues regarding MoBIE, please open an issue on github (https://github.com/mobie/mobie-viewer-fiji) or create a topic on image.sc with the tag “mobie”.

### State of the art / existing tools

Visualizing, exploring and sharing massive multi-modal data poses significant challenges. The size of the data can be too large to fully load into a computer’s memory, too large to download in a reasonable amount of time or even too large to store on a regular hard drive. These challenges can be addressed by loading only the currently shown image data on demand. To support this use-case, next-generation file formats such as OME-Zarr^1^ and N5^2^ have been developed. They store multi-dimensional (pyramidal multi-scale) image data in chunks (small data blocks), and store each chunk in a separate file. This design enables chunk level access of the image data in object storage, which is commonly used to store and distribute big data online.

Several performant, webGL based, image viewers^3–6^ can read next generation file formats. Most of them currently require setting up a dedicated server to access the image data (webKnossos^6^, Neuroglancer^5^, Catmaid^3^), only Viv^4^ currently runs purely on the client side and does not need a server. Of these tools, Neuroglancer is most similar to MoBIE, sharing several functionalities such as arbitrary slicing of 3D data, interactive rendering of segmentation data and a JSON based schema to specify the viewer state. In the future, we plan to enable accessing the same data in MoBIE and Neuroglancer through shared support of OME-Zarr and development of a compatible JSON metadata standard. Many of these web applications require external data conversion to their supported formats, as well as, for most of them, setting up a web server. The need for this infrastructure makes them hard to use for biologists without significant computational expertise. Furthermore, most are stand alone tools and not directly integrated into one of the commonly used bioimage analysis platforms (see below), making their integration with well established workflows for common analysis tasks challenging.

Many biologists use a bioimage analysis platform for their data visualization and analysis needs. Popular platforms include Fiji^7^ and Icy^8^, which are Java based, as well as napari^9^, which is Python based. Among those tools, Fiji stood out for our purposes as it is very widely used, and is integrated with ImgLib2^10^, a general purpose n-dimensional image processing Java library that was designed for big image data and offers performant lazy evaluation and caching implementations. ImgLib2 also provides the primitives needed for n-dimensional image registration, which are used in the BigWarp^11^ and BigStitcher^12^ plugins. Furthermore, Fiji includes the ImgLib2-based BigDataViewer^13^ (BDV), which provides performant slice viewing of (TB-sized) multi-resolution 3D time-lapse data, as well as Bio-Formats^14^, a Java library for translating image data from over 160 proprietary file formats using standardized, open formats. In summary, Fiji is widely used in the bioimage community, offers diverse bioimage analysis functionality, and contains performant core libraries for big image data visualization and registration. Thus, we decided to implement MoBIE as a Fiji plugin. See below section on MoBIE Viewer Core Features and MoBIE viewer Implementation for more details.

### MoBIE Plugin usage

MoBIE can be installed in Fiji by enabling the “MoBIE” update site. It is then available via “Plugins->MoBIE” (or the plugin search), where several commands for opening and creating projects are implemented. The most common way to open a project is via “Plugins->MoBIE->Open->Open MoBIE Project…” and entering the project’s address, for example https://github.com/mobie/platybrowser-project. Upon loading, MoBIE will display a predefined default view. Other views can be selected from the main MoBIE menu. Under “Plugins->MoBIE->Open->Advanced” three commands that offer specifying advanced options for loading projects can be found, including “Open MoBIE Project with S3 credentials.”, which allows users to enter the credentials to open projects that contain data stored on non-public S3 storage. Furthermore, the “Open” menu contains two commands for convenient access to special MoBIE projects: “Open Published MoBIE Project …” provides direct access to a list of published projects and “Open MoBIE Published View …” to a list of some of the figures within these projects. For example, the views corresponding to panels a, b and d of Fig. 1 are available as “MoBIE: Figure 1a”, etc. These views are of course also accessible from the corresponding projects. Furthermore, “Plugins->MoBIE->Create” contains the commands for creating and editing MoBIE projects with an interactive user interface.

### MoBIE viewer core features

MoBIE is built using BigDataViewer^13^ (BDV) and ImgLib2^10^. To facilitate readability we describe the core features of MoBIE without making a distinction of whether the implementation was mainly done by us or whether the functionality existed already in the libraries that we rely on. To check for our contributions please see the section on Additions to the current BDV/ImgLib2 ecosystem.

#### Image visualization

For the interactive exploration of image data, MoBIE relies on BigDataViewer (BDV), which provides arbitrary plane slicing of TB-sized, chunked and pyramidal 3D time-lapse image data. MoBIE supports the chunked pyramidal image data formats OME-Zarr, BDV-N5 (local and remote access) and BDV-HDF5 (local access). See “Project structure, storage and metadata” for details.

MoBIE provides a user interface for managing the display of many images. The user interface allows grouping *views*, where each *view* corresponds to a full configuration of the viewer, into several drop-down menus. One view can represent more than one image and, upon selection, will automatically configure a variety of display settings (see below for more details). It can contain one or multiple *image displays*, which each contain one or multiple images that share the same rendering settings (color, contrast limits, opacity). Each image display is added to the display adjustment panel, which contains buttons to remove all corresponding images (“X”), toggle their visibility (“S”), or adjust their contrast limits (“B”) and color (“C”) (see Fig. 1a) for a screenshot of this menu). To efficiently remove all images from the current display we provide a “clear” button. When many images are shown simultaneously it can become difficult to know what’s what. To alleviate this challenge we provide an “overlay names” checkbox, which will trigger the rendering of the image names within BDV, next to the respective images.

In addition, MoBIE implements an alpha-blending rendering mode in which the opacity of images can be changed via the “O” button. Depending on the position of the images in the display adjustment panel and the opacity setting, images can occlude other images that are higher up in the list. This is for example useful to combine multiple acquisitions of the same object such that the most informative (e.g. highest resolution) version is visible, occluding less informative (e.g. lower resolution) images of the same object (see Fig. 1d). Such alpha-blending is also useful to check image registrations as by moving the opacity slider some images (against which one may have registered) become more or less visible. For efficient navigation in large spatially distributed image datasets, MoBIE provides a focus (“F”) button that centers the current view onto the respective image(s). There are more ways to navigate between images as described in the below paragraphs on tables and scatter plots.

The “V” checkbox supports volume rendering of images and segmentation objects (s.b.) using the Fiji 3D Viewer^15^. MoBIE also adds a right-click context menu to the BDV window. If applicable, context menu items will only operate on images currently visible at the location of the mouse pointer. The “Take Screenshot” menu item renders the current view as a 2D image in the traditional ImageJ viewer where it can be further modified and saved. The sampling of the rendering can be freely adapted, e.g., for creating images with high spatial detail and a specific pixel size. Renderings are done both as colored RGB and multi-channel gray scale images. The “Show Raw Image(s)” menu item provides access to the pixel raw data at one of the available resolution levels. The “Configure Image Volume Rendering” menu item allows one to specify the 3D rendering resolution. This feature is not compatible with big image data as for rendering all data at the requested resolution level must be loaded into memory; we thus recommend conservative usage of this functionality. There are several more context menu items that will be described in the corresponding paragraphs below.

#### Image transformations and registration

The visualization of multi-modal image data requires the 3D placement and scaling of images of different dimensions (number of voxels) in data space in a common physical coordinate system (physical space). The BDV image data model implements this by attaching an affine transformation that maps data space to physical space to each image. Leveraging this core functionality, MoBIE provides a number of additional image transformation modes. It allows the presentation of selected sub-volumes or areas of a source as cropped detail views.

MoBIE also allows specifying an automated arrangement of images in nested 2-D grids, including images to which another transformation (e.g. crop) has been applied already. This is useful, e.g., for the visualization of high-throughput LM (see Fig. 1b) and EM data (see Fig. 1d) when comparing several images of similar features with each other that originally are located at different positions.

For image registration, BDV supports the interactive manual alteration of an image’s affine transform. In addition to this functionality, MoBIE offers a convenient integration of the BigWarp Fiji plugin^11^, which enables the registration of images via landmarks. See Supp. Video 3 for a demonstration of both registration features.

#### Viewer transformations

The image plane rendered by BDV is defined by the “viewer transformation”, an affine transformation that maps the BDV window into the viewer’s global coordinate system (physical space). Leveraging this feature, MoBIE provides several means of specifying the rendering location. In addition to the standard affine transformation MoBIE supports a “normalized affine transformation”, which does not depend on the size of the BDV window and thereby works better for sharing the viewer state among different users. In addition, one can specify a “normal vector transformation” where MoBIE will align the current image plane perpendicular to the given vector. This is useful for navigating in an anatomical coordinate system, for example given by the dorsal/ventral axis of an organism. Finally, one can specify a “position transformation”, where a 3D location will be centered, without adjusting rotation or scale. This can be used to locate points of interest, for example specific cells or organs in a dataset showing an animal. All viewer transformations also include the time point at which the images should be shown.

#### Segmentation image data visualization

In addition to the visualization of intensity images, an important feature of MoBIE is to provide comprehensive functionality for the exploration of segmentation images. Technically, these images are represented as label mask images (2D or 3D and time), where pixel values encode object indices. The default visualization is a lookup table where different pixel values are assigned random but well discernible colors^16^. A common issue with such a visualization is that neighboring objects may by chance have very similar colors that are hard to distinguish. MoBIE thus provides both a context menu item (“Change Random Coloring Seed”) and a keyboard shortcut {CTRL + L} to shuffle the random colors. The label rendering can be further configured via the context menu item “Configure Label Rendering”. Labels can be either rendered covering the whole object or only its boundary. Image segments can also be colored based on measured features, stored in an image segment feature table (s.b.). One can specify any column in the table and choose from a number of color maps (both coloring of numeric or categorical values is supported). MoBIE also supports selecting (multiple) image segments by {Ctrl + Click}; upon selection the coloring mode changes such that selected segments are standing out.

Selected image segments can be rendered in 3D by activating the “V” checkbox in the display adjustment panel. For volume rendering MoBIE employs Fiji’s 3D Viewer^15^ and syncs the coloring of the segments with the coloring in BDV. 3D meshes are computed on the fly and their level of detail can be adjusted in the “Configure Segments Volume Rendering” context menu item. Selecting a new segment for 3D visualization in the BDV window will automatically center the 3D viewer onto the new segment; vice versa, clicking on a segment in the 3D viewer will center the BDV view onto that segment.

#### Tables (Annotations)

A core feature of MoBIE is an interactive table display, which is critical for the efficient exploration of large image datasets. Table rows either correspond to image segments (s. a.) or regions (s. b.). The coloring and selection of image segments or regions are synched with the image display in BDV. Also navigation is synched as clicking on a table row will center the image display on the respective segment or region; vice versa, selecting an image segment or region in the image display will move the corresponding table row to the top of the current table display. Tables can be sorted by clicking on the column headers. Tables have a menu with items: “Table”, “Select”, “Color” and “Annotate”. The “Table” menu provides methods to save the table or load additional columns that will be merged into the current table. The “Select” menu provides functionality to select a subset of rows based on the values in any column. Table rows (and the corresponding image segments or regions) can be colored corresponding to the values in any column choosing from a number of color maps (“Color > Color by Column.”): random colors (categorical data), heatmaps (numerical data), or fixed color schemes (ARGB columns). The “Annotate” menu provides functionality to manually annotate image segments or regions. The corresponding annotations will be stored as a new column or can be appended to an already existing column. Upon selecting either “Create New Annotation” or “Continue Annotation” a user interface will appear that enables annotating one or several objects as belonging to one of several freely configurable classes. The selection can be done via {Ctrl + Click} in the table, BigDataViewer or scatter plot window or using the “Select” table menu (s.a.). For efficient annotation of subsequent objects, “Select next” and “Select previous” buttons are provided. For efficient selection of a specific object in a large dataset it also is possible to select an object by its identifier (“label_id” or “region_id”). Annotations are added to the respective annotation table column, which can be saved and loaded.

#### Scatterplots

As an additional means for the exploration of image segment or region features, MoBIE provides an interactive scatter plot, where two table columns are plotted against each other. Coloring and selections are synched with the image and table displays. Via the “Configure point(s) selection” context menu item one can specify a radius for the selection of points in the scatterplot around the mouse cursor position. This is, e.g., useful for the efficient selection of multiple image segments with similar morphological or other measured features.

#### Image region visualization

MoBIE supports the annotation of regions comprising one or multiple images, e.g. a well in a high-throughput microscopy experiment. Each region is rendered as the union of all contained images and associated with a table row. The region visualization features are essentially identical to the ones described above in the sections about image segmentation and table visualization (s. a.). In brief, regions can be consistently colored, selected, and navigated in the image, table and scatter-plot displays.

#### Views

An important core feature of MoBIE is an expressive JSON schema for specifying complex views on many images, see “Project structure, storage and metadata” for details. Views can save the full viewer configuration. They can be saved via the context menu and loaded again by selection of the view name from a drop-down menu in the main MoBIE menu. Each MoBIE dataset can specify an arbitrary number of views that are available by default and additional views can be loaded from JSON files via the context menu. The view features enable seamless configuration of any MoBIE configuration by arranging it in MoBIE and then saving the view. These saved views can then be shared with other MoBIE users, for example to point out locations in the data or full analysis results that underpin a research result. They can also be used to configure interactive figure panels, such as the figure panels in this publication. Views can also be created programmatically with our python library, see Project creation for details.

### MoBIE viewer implementation

The MoBIE viewer is implemented in Java. The majority of the source code is contained in two github repositories: https://github.com/mobie/mobie-viewer-fiji and https://github.com/mobie/mobie-io. The *mobie-io* repository contains the core functionality for the loading and writing of various big image data formats (BDV-HDF5^13^, BDV-N5^2^, OME-Zarr^1^) from and to file systems or S3 object stores. The *mobie-viewer-fiji* repository implements the Fiji plugins for big image data loading, writing and visualization (see above section on MoBIE’s core features). The code builds upon various other java libraries, most notably ImgLib2^10^, BigDataViewer^13^, ImageJ 3D-viewer^15^, N5^2^ and bigdataviewer-playground (https://github.com/bigdataviewer/bigdataviewer-playground).

### Project structure, storage and metadata

In order to be accessible via the MoBIE plugin, data has to be organized according to the *MoBIE Project specification*. A project may contain several *datasets*, which contain the image data, tabular data and view metadata that can be displayed by a single MoBIE instance. Different datasets in a project could for example correspond to separate specimens from an experiment or to versioned analysis results of the same data. Each dataset corresponds to a sub-folder in the root project directory, which also contains a *project.json* file that lists all available datasets and contains some further project metadata. This metadata includes the image file formats used in the project, which determine the available modes of data access.

Project data can be accessed in different ways, depending on the type of data: image data can be accessed either from the local filesystem or from object storage. Tabular data and metadata can be accessed from the local filesystem, object storage or github. This allows three different access modes for a project: i) all data is read from the local filesystem, ii) all data is read from object storage, or iii) image data is read from object storage and tabular data as well as metadata is read from github. Options ii) and iii) enable data access in the cloud, and thus enable convenient sharing of MoBIE projects with other researchers. Data can be made fully public, using non-access-restricted object storage and github repositories, or in a closed collaboration, using object storage with credential based access right management. In order to access image data in object storage we rely on modern file formats that allow on-demand chunk access, see next paragraph for detail. Currently, we only support remote data access in object storage with an Amazon S3 compatible API; but extension to other protocols such as Azure Blob Storage would be possible. Note that the S3 API is open source and tools such as min.io (https://min.io/) that allow setting up on-premise S3 object storage exist. Consequently reliance on the S3 API does not result in vendor lock-in.

All metadata associated with MoBIE is stored in JSON files, which follow a json schema (https://json-schema.org/), a JSON based language to describe metadata formats. This schema can be used to statically verify the correctness of any MoBIE project. Specifically, we define schemas for the project.json file (see first paragraph), the dataset.json file (see next paragraph) and JSON files for view specification. The MoBIE schemas are available at https://github.com/mobie/mobie.github.io/tree/master/schema.

A dataset contains the data available to a MoBIE plugin instance; datasets can be switched using the “dataset” dropdown in the plugin’s main menu. Each dataset has a root level folder that contains the *dataset.json* file, which specifies the sources and views available. Sources correspond to image data. We currently support two different types of sources: *imageSource* for intensity images and *segmentationSource* for label mask images. For both source types the metadata specifies image data formats and data locations. We currently support the following formats: OME-ZARR^1^, BDV-N5 (https://github.com/bigdataviewer/bigdataviewer-core/blob/master/BDV%20N5%20format.md) and BDV-HDF5^13^. Both BDV file formats use an additional xml file that specifies image metadata; OME-ZARR stores this metadata directly in the zarr file that contains the image data. For local data access the relative path to the xml or zarr files is stored in the metadata. For remote data access via object storage, the full address of the xml / zarr file on S3 is stored. Only OME-ZARR and BDV-N5 support remote data access, BDV-HDF5 can only be accessed locally. We also offer experimental support for the n5 based OPEN-ORGANELLE data format, see also Access of data deposition archives. Below table shows the supported read and write options for each file format.

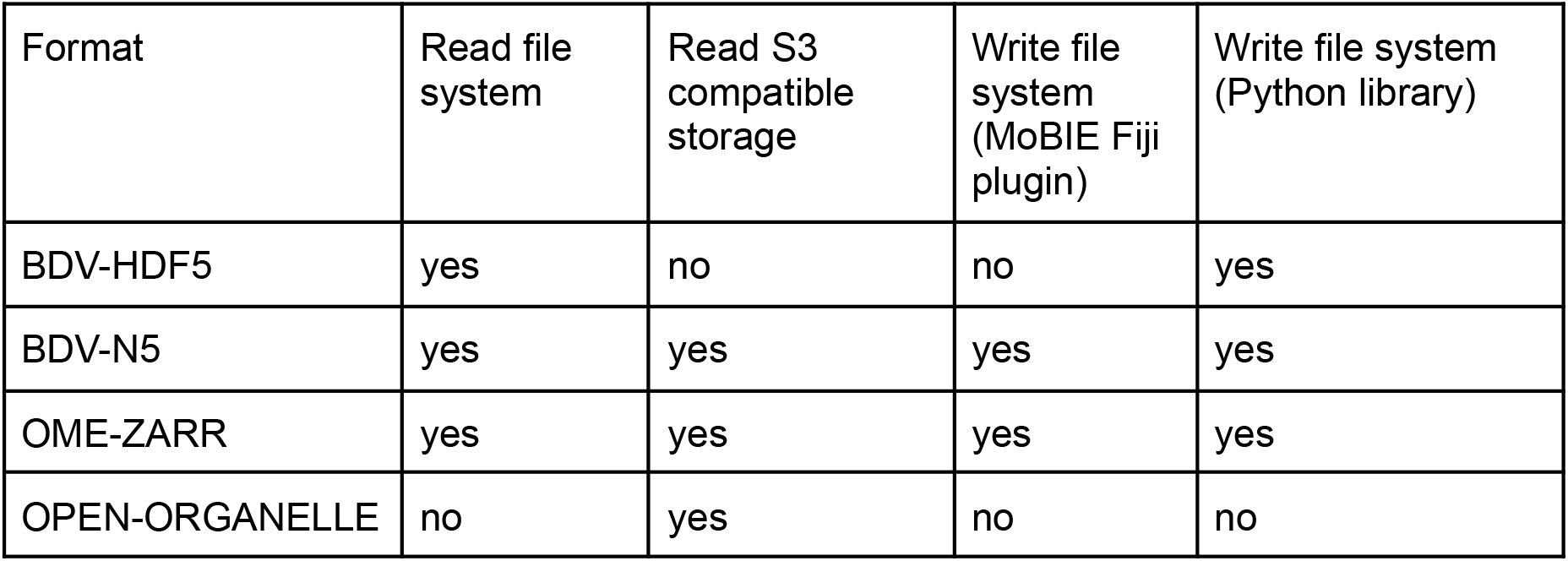

| Format | Read file system | Read S3 compatible storage | Write file system (MoBIE Fiji plugin) | Write file system (Python library) |
| --- | --- | --- | --- | --- |
| BDV-HDF5 | yes | no | no | yes |
| BDV-N5 | yes | yes | yes | yes |
| OME-ZARR | yes | yes | yes | yes |
| OPEN-ORGANELLE | no | yes | no | no |

Segmentation sources can additionally specify the location of tabular data stored as tab separated values (tsv). For each segmentation the relative path to a folder, which contains arbitrarily many tables can be given. These tables must contain a column called *label_id*, which maps to the label mask ids in the corresponding image data, and may contain arbitrary additional columns. By convention, image data should be stored in a subfolder called *images*, which should contain subfolders for each image format. Tables should be stored in a subfolder called *tables*, with a separate subfolder for each segmentation (or region display, see below) with associated tabular data. Datasets may contain another subfolder called *misc* for other data, such as JSON files specifying additional views (see below). See the image to the right for the recommended MoBIE project folder structure.

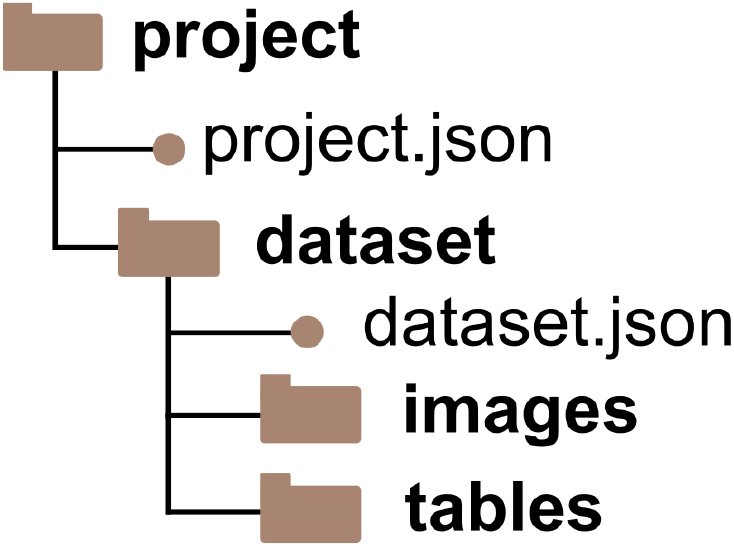

In addition, the dataset metadata contains a list of views, which correspond to pre-set viewer configurations that are selectable in the MoBIE menu. A view consists of the following elements: *sourceDisplays* define the image or segmentation sources loaded in the view, and their display settings such as brightness, opacity or blending modes. For segmentation sources they also contain the tables that will be loaded and visible as well as selected segments, if any. In addition, they may also contain *regionDisplays*, which enable annotations for specific areas in the viewer coordinate space, e.g. annotations for each position in a grid transformation. The region displays must have an associated table, which has a mandatory column *region_id*, and otherwise share the same properties as the segmentation tables. Next, *sourceTransforms* define the on-the-fly transformations, such as affine, grid or crop transformations, that are applied to the sources in the view. Finally, the *viewerTransform* defines the “camera position” of the view.

A full description of the MoBIE project specification, including examples for the JSON specifications, is available at https://mobie.github.io/specs/mobie_spec.html, the version of the specification at time of writing is available at https://mobie.github.io/v021/.

### Project creation

#### Fiji

The MoBIE Fiji plugin includes a series of tools to simplify the creation of MoBIE projects. This project creator is implemented in Java, with the source code included in the main MoBIE repository: https://github.com/mobie/mobie-viewer-fiji. It can convert images that are loaded into Fiji to both the BDV-N5 and OME-Zarr file formats using functions from the *mobie-io* repository: https://github.com/mobie/mobie-io. This conversion builds on top of the export functions from the *bigdataviewer_fiji* (https://github.com/bigdataviewer/bigdataviewer_fiji), *bigdataviewer-core* (https://github.com/bigdataviewer/bigdataviewer-core) and Saalfeld lab *n5-zarr* (https://github.com/saalfeldlab/n5-zarr) repositories, adding various new features including downsampling methods for label mask images.

The project creator provides a graphical user interface, allowing easy creation of projects with no programming experience. Datasets can be created by clicking the ‘Add’ button on the dataset row, and edited by clicking the ‘Edit’ button. Images can be added to a project by opening them in Fiji and then clicking the ‘Add’ button on the image row. This allows various metadata to be specified including the image name, the MoBIE dropdown menu it should appear in, and an affine transform. Images are then automatically converted to BDV-N5/OME-Zarr with all the required metadata for MoBIE. Images that are already in a compatible file format can also be added to the project by selecting the ‘bdv format image’ option after clicking the ‘Add’ button. Once a project is complete, it can be opened directly in MoBIE by clicking the ‘Open in MoBIE’ button.

The project creator also supports adding the required metadata for a remotely hosted project. Once this is complete, the entire project can be uploaded to S3 and accessed from anywhere in the world by opening the S3 location in MoBIE. The project creator can also be scripted with any of the Fiji scripting languages, allowing easy creation of larger / more complex projects. Detailed use instructions are available in the ‘Making your own projects’ section of the MoBIE documentation at https://mobie.github.io/ and Supp. Video 5 shows a live demonstration of the project creation process.

#### Python

We also provide a python library for creating, extending and validating MoBIE projects. The library is available at https://github.com/mobie/mobie-utils-python. It uses https://github.com/constantinpape/cluster_tools to enable conversion of big data into the file formats supported by MoBIE and generation of segmentation tables. It supports execution on the local machine or on a cluster computer using slurm or LSF scheduling systems. Furthermore, the library implements functions for generating any view configuration supported by MoBIE. It also implements validation functions to check whether a given project, dataset or view file adheres to the MoBIE specification. An example notebook that demonstrates the usage of *mobie-utils-python* is available at https://github.com/mobie/mobie-utils-python/blob/master/examples/create_mobie_project.ipynb.

### Additions to the current BDV/ImgLib2 ecosystem

As mentioned, the MoBIE viewer builds on ImgLib2, BigDataViewer and N5-java. Here, we will briefly list our additions in terms of code and functionalities that, to the best of our knowledge, are not available in the current ImgLib2, BigDataViewer and N5 libraries. We have in the past closely collaborated with their developers. Striving for a coherent eco-system, we will work with these developers to choose and contribute MoBIE features to Imglib2, BigDataViewer, N5-java or other appropriate general purpose libraries.

- Coloring of label mask (segmentation) images in BDV with various color maps and based on label feature (measurements or annotations) values
- Configurable boundary rendering mode for label mask images.
- Interactive selection of labels in BDV via {Ctrl + Click}
- On-the-fly mesh computation and volume rendering of selected labels (this feature is also available in Paintera, a standalone BDV based application, but not in the standard BDV)
- Interactive feature table, coloring and selection synchronized with label mask visualization in BDV, including {Ctrl + Click} selection
- Interactive scatter plot of table columns, coloring and selection linked with table display and BDV, including {Ctrl + Click} selection
- Right-click context menu in BDV
- Screenshot functionality (available via context menu) for BDV with adjustable sampling, e.g. for creating high quality figures at a defined resolution (BDV supports movie recording at the current screen resolution)
- Show raw data functionality for BDV for opening a selectable resolution level of any shown image in the standard Fiji viewer
- Reading of unsigned long data type for HDF5/XML images, as needed for label mask images with many labels (BDV supports unsigned short data type, limiting the maximal number of labels to 65535)
- “Center” and “mode” downsampling methods for writing multi-scale label mask images (BDV provides “average” downsampling, which is not suited for label masks)
- Reading (local and remote) and writing (local) the latest versions of OME-Zarr
- User interface for manual annotation of table columns, corresponding for segmentation or region tables.
- Region annotations where groups of images can be overlaid with a color based on feature values

○ Essentially all functionality available for segmentations (label masks) are also available for regions (e.g. color maps, selection, tables, scatterplot)
○ Internally, regions are computed on the fly based on the position of the annotated images, thus, in contrast to segmentations, no precomputed images need to exist on disk
- Alpha-blending mode for BDV, e.g., to overlay high-resolution EM data onto low-resolution data with varying occlusion (“O” button)
- Focus button (“F”) to center BDV on the respective image (important to navigate in large image data sets)
- JSON schema to read and write the MoBIE viewer state, including image display settings, label selections, table and scatter plot visibility, 3D renderings, and various image transformations (serialization of affine transformations were already supported by BDV in XML format).
- BigWarp integration for any pair of images currently visible in BDV
- Crop transformation to only render parts for an image
- Normalized affine transformation to render a similar view independent of the BDV window size
- Nested grid transformation to automatically lay out images in a 2D grid
- Project creator plugin to create MoBIE projects using the GUI

### Example MoBIE projects

Building a high-resolution cellular atlas for *Platynereis dumerilii* for Vergara et al.^17^ has been our initial motivation for developing MoBIE. The atlas contains a complete 6 day old larva imaged in EM, corresponding to 8 Terabytes of raw data, and spatial expression maps for about 200 genes, that were obtained by averaging LM volumes of fluorescence in-situ hybridisation for several animals of the same developmental stage. It also contains the segmentations of all cells, nuclei and selected organelles, derived from the EM data. The gene expression maps are registered to the EM coordinate space by matching the signal from their nucleus channel with the nucleus segmentation. Finally, the atlas contains several feature descriptors and cluster assignments for the cells, which are derived from the cellular morphology and the gene expression. In Fig. 1a we show a cluster assignment mapped to the cell segmentation in the image space (ii) and a low dimensional representation of the gene expression space (iv) as well as all cells belonging to one of the clusters rendered in 3d (v).

MoBIE enables parallel browsing of registered multi-scale data as resulting from correlated light and electron microscopy (CLEM) at native resolutions without the need of data interpolation. We visualize an entire CLEM experiment acquiring two and three-dimensional EM data and 2D fluorescence data on resin sections^18^ containing genetically labeled HeLa cells^19^ registered in its spatial context in Fig. 1d. Sources with bright background such as brightfield or EM images and those with dark background such as fluorescence data are combined using MoBIE’s alpha and additive compositing modes. We also employ its annotation features to highlight the bounding boxes and positions of the individual sources for the different figure panels. The individual sources were registered on top of each other using the transformation tools available in MoBIE. We can also visualize the high-resolution content of the dataset in parallel using a grid view to enable direct visual comparison of morphological features.

We demonstrate MoBIE’s utility for high-throughput microscopy (HTM) on data from an immunofluorescence assay for SARS-CoV-2 antibodies^20^. Briefly, the assay uses Vero E6 cells that are partially infected with the virus, mixes the cells with human serum and images them on a 96 well plate. Three channels are acquired, staining nuclei, viral RNA and IgG antibodies in the serum. To determine the antibody response all individual cells are segmented based on the nucleus and serum signals. The cells are then classified into infected and non-infected cells based on the virus marker expression per cell, and the antibody response is measured through the ratio of serum intensity between infected and non-infected cells, with larger ratios corresponding to a stronger response. Fig. 1b shows data from this assay as well as analysis results in MoBIE: a) shows the nucleus channel (blue) and serum channel (green) for the whole plate. The plate layout is created using a nested grid transformation. b) shows the measured antibody response scores per well, using the table coloring functionality. c) shows a zoom-in for one of the wells, which can be conveniently accessed via the corresponding row in the well table. It also showcases the table annotation mode that is used here to label outlier images, which should be excluded from the analysis. d) shows the cell segmentation for one of the images from c) and uses the table coloring functionality to display which cells were classified as infected and which as non-infected.

We have used MoBIE to deliver advanced visualization for several more projects beside the three examples highlighted here, an overview of additional projects is shown in Suppl. Fig. 1. These projects include cells infected with SARS-COV-2 and imaged with EM tomography^21^ (c), where MoBIE’s grid arrangement is used to show the tomograms in a gallery and the corresponding image table contains organelle annotations for the individual tomograms. In the same study SARS-COV-2 infected cells were also imaged in volume EM, shown in (d), with additional organelle segmentations. Next, (e) shows a time series of a developing *Arabidopsis thaliana* root imaged with lightsheet microscopy^22^, demonstrating MoBIE’s time-lapse support for both intensity image and corresponding segmentation data. See Supp. Video 2 for details. The panel (f) shows zebrafish embryos imaged with lightsheet microscopy^23^, where MoBIE’s grid transform is used to arrange the individual volumes in a gallery. In addition, the cell segmentations for all volumes are shown and the segmentation table contains morphological measurements for the cells in all volumes. Next, (g) shows plankton cells imaged in EM^24^, including organelle segmentations. Panel (h) shows a sponge choanocyte chamber imaged in EM^25^ and zooms in on the spatial interaction of a neuroid cell with several choanocytes, which is also rendered in the 3d viewer. Finally, (j) shows neural tissue imaged in EM from a part of the mouse brain^26^.

### Access of data deposition archives

Increasingly, imaging data from publication in the life sciences is deposited in online image archives to ensure reproducibility of the corresponding study and enable re-use of the data for other research purposes. Many journals and funding agreements also mandate this data deposition. In most cases, this data is available only for bulk download, which makes re-use of big datasets challenging since a potential user would first need to download a large amount of data before assessing if it even fits their needs. On demand access of image data in online deposition archives is thus highly desirable to enable interested researchers to quickly check this data, or even enable them to perform full data exploration and analysis on demand. With its support for many data modalities, extensive features for data exploration and analysis and capability to stream image and tabular data on demand, MoBIE is an ideal candidate application for this task. In fact, we have already implemented prototypes for on demand access of data from two online deposition archives.

The first is Open Organelle^27^, which is a data deposition archive hosted by HHMI Janelia Research Campus for storing very high resolution EM volumes of cells, derived organelle segmentations and correlative LM data. It uses n5 to store the data, which is publicly accessible via S3 storage and relies on a custom image metadata format. We have added experimental support for this image data format to *mobie-io*, and can thus create MoBIE projects that access data via Open Organelle. A test project demonstrating this functionality is available at https://github.com/mobie/open-organelle-test. It uses the “jrc-hela-2” cell from the openOrganelle cell repository and only contains the EM data and the endosome segmentation. Adding further segmentations would be simple. Creating and extending Open Organelle compatible projects is also supported by the MoBIE python library.

The second is the BioImage Archive^28^, an image data repository hosted by the European Bioinformatics Institute. It accepts image data submissions associated with publications in the life-sciences. It stores this data on S3 compatible object storage, so data deposited in a format supported by MoBIE can be streamed from it. We demonstrate this capability with an example project that corresponds to a subset of the data from Cortese et al.^21^. It is available at https://uk1s3.embassy.ebi.ac.uk/bia-mobie-test.

## Supplementary Material

### Online documentation

- Supp. Note 1: The current version of the MoBIE documentation is available at https://mobie.github.io/. The version corresponding to the state of the manuscript submission is available at https://mobie.github.io/v021/.

### Online videos

These are only available on YouTube, since upload through the journal submission system failed because of the file size.

- Supp. Video 1: Detailed MoBIE functionality in the platybrowser: https://youtu.be/oXOXkWyIIOk
- Supp. Video 2: Time series project: https://youtu.be/Md4PbK50NE0
- Supp. Video 3: Registration in MoBIE via manual moving of sources and/or BigWarp: https://youtu.be/jKlM68lrhso
- Supp. Video 4: Object annotations: https://youtu.be/M-QUE-Qh97w
- Supp. Video 5: Fiji project creation: https://youtu.be/3oP3t6elsQU

### Supplementary figures

**Supplementary Figure 1:**
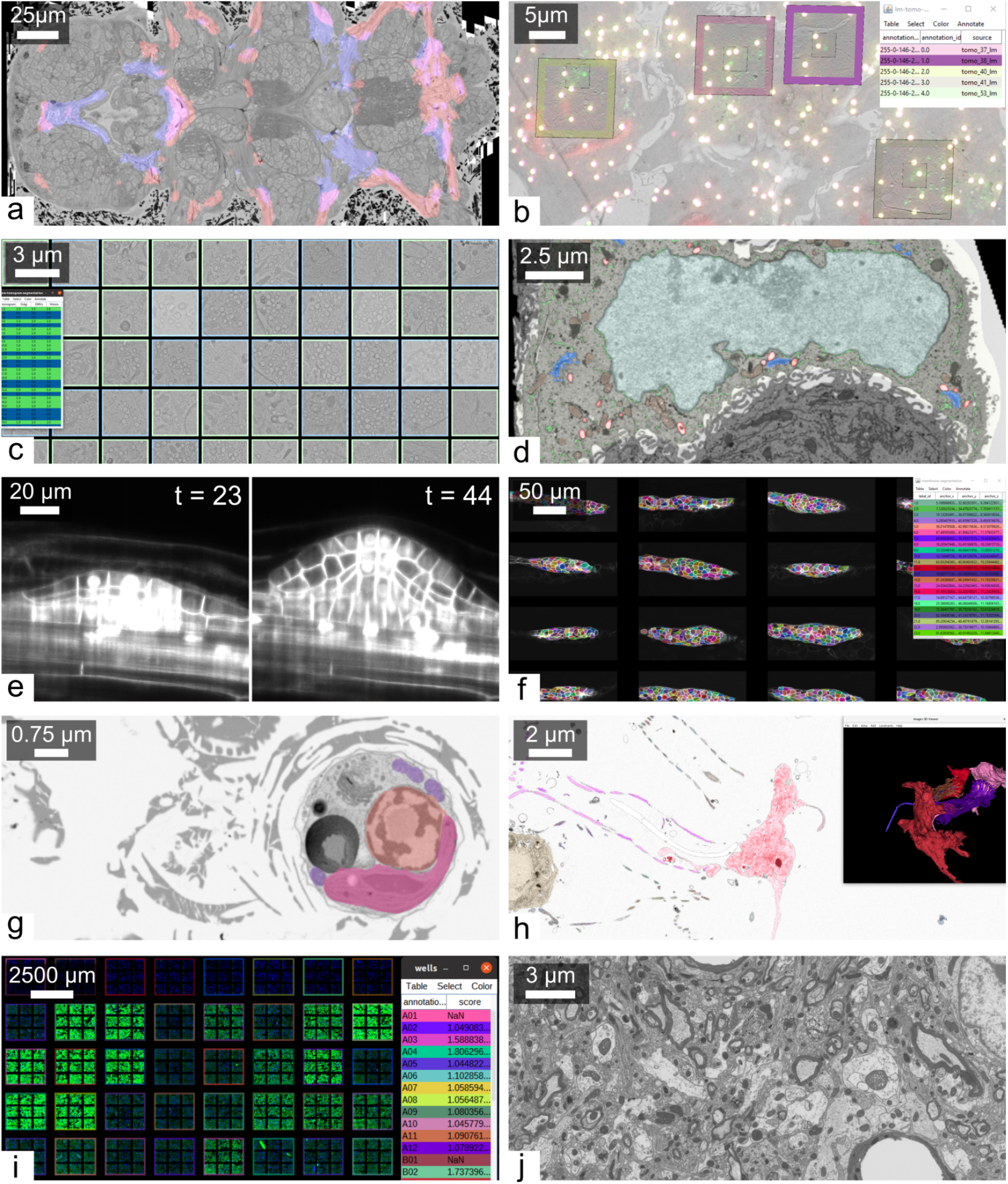
Overview of MoBIE projects - **a** cellular atlas of *Platynereis dumerilii* made up of whole organism EM of a larva^17^, gene expression maps registered to the EM (two gene expression maps shown in red and blue) and cell segmentation (not shown), see also Fig. 1a. **b** CLEM project with overview EM slice, fluorescence signal as well as low and high resolution EM tomograms, see also Fig. 1d. **c** grid arrangement of EM tomograms of SARS-COV-2 infected cells^21^, the site table contains information about the organelles present in each tomogram. **d** volume EM of SARS-COV-2 infected cells^21^, including segmentation of organelles. **e** lightsheet time-series of developing *Arabidopsis thaliana* roots^22^, see also Supp. Video 2. **f** lightsheet volumes of zebrafish embryos^23^ with cell segmentation and derived morphological measures in grid arrangement. **g** plankton cells imaged in EM^24^ with organelle segmentations. **h** cells in a sponge choanocyte chamber imaged in EM^25^, with segmented cells and microvilli, which are also shown in the 3d viewer. **i** immunofluorescence assay for SARS-COV2 antibodies^20^, imaged with high-throughput microscopy and shown in the multi-well plate layout recreaded with two nested grid transformations. **j** EM volume of neural tissue from a mouse brain^26^.

